# Machine Learning Prediction of Non-Coding Variant Impact in Human Retinal *Cis*-Regulatory Elements

**DOI:** 10.1101/2021.10.19.464837

**Authors:** Leah S. VandenBosch, Kelsey Luu, Andrew E. Timms, Shriya Challam, Yue Wu, Aaron Y. Lee, Timothy J. Cherry

## Abstract

**Purpose:** Prior studies demonstrate the significance of specific *cis*-regulatory variants in retinal disease, however determining the functional impact of regulatory variants remains a major challenge. In this study, we utilize a machine learning approach, trained on epigenomic data from the adult human retina, to systematically quantify the predicted impact of *cis*-regulatory variants.

**Methods:** We used human retinal DNA accessibility data (ATAC-seq) to determine a set of 18.9k high-confidence putative *cis*-regulatory elements. 80% of these elements were used to train a machine learning model utilizing a gapped k-mer support vector machine-based approach. *In silico* saturation mutagenesis and variant scoring was applied to predict the functional impact of all potential single nucleotide variants within *cis*-regulatory elements. Impact scores were tested in a 20% hold-out dataset and compared to allele population frequency, phylogenetic conservation, transcription factor (TF) binding motifs, and existing massively parallel reporter assay (MPRA) data.

**Results:** We generated a model that distinguishes between human retinal regulatory elements and negative test sequences with 95% accuracy. Among a hold-out test set of 3.7k human retinal CREs, all possible single nucleotide variants (SNVs) were scored. Variants with negative impact scores correlated with reduced population allele frequency, higher phylogenetic conservation of the reference allele, disruption of predicted TF binding motifs, and massively-parallel reporter expression.

**Conclusions:** We demonstrated the utility of human retinal epigenomic data to train a machine learning model for the purpose of predicting the impact of non-coding regulatory sequence variants. Our model accurately scored sequences and predicted putative transcription factor binding motifs. This approach has the potential to expedite the characterization of pathogenic non-coding sequence variants in the context of unexplained retinal disease.

## Introduction

Retinal disorders affect over 2 million individuals worldwide and consist of many classes of disease. Over 260 genes have now been associated with retinal disorders.^1,2^ However, as many as half of all cases cannot be explained by variants in protein-coding genes alone.^3^ This suggests that risk variants located within the non-coding genome may contribute to retinal disease. The comparatively vast non-coding genome harbors *cis*-regulatory elements (CREs) including promoters, enhancers, silencers, and boundary elements that play a critical role in gene expression^4-7^. Genome-wide association studies (GWAS) frequently link non-coding regions to disease phenotypes ^6,8-11^. Moreover, individual case studies have identified causal regulatory variants in retinal disorders including Blue Cone Monochromacy, Non-syndromic Congenital Retinal Non-Attachment, and Aniridia with Foveal Hypoplasia^12-14^. However, due to the poor functional characterization of non-coding regions, it remains a challenge to systematically interpret the impact of variants within CREs.

CRE function is mediated by complex interactions between transcription factors (TF) and DNA sequences^15,16^ to yield the appropriate transcriptional profile for a given cell type.^17^ These interactions can be characterized through assays for DNA accessibility (ATAC-seq and DNase-Seq) and protein binding (ChIP-Seq, CUT&RUN, and CUT&Tag) to identify and characterize candidate CREs in a given tissue or cell type at a single point in time.^18,19^ Despite recent advancements, it remains challenging to understand the functional significance of genetic variants within CREs without further experimental or integrative computational analyses.^20,21^ Identifying and investigating all potential regulatory regions and putative variants is a monumental task that requires painstaking efforts.

Recent developments in artificial intelligence have popularized the use of machine learning for the holistic interpretation of multi-modal epigenetic sequencing data.^22,23^ Many different approaches have been developed to accurately predict the inferred value of genetic sequences including non-coding regulatory regions.^24-26^ Such approaches have demonstrated promise in select cell lines and tissue types, and have been used successfully to integrate epigenomic data in the context of the human retina.^27^ This supports the premise of a comprehensive, tissue-specific analysis for CRE variant prioritization in the human retina. While a number of approaches are available to predict sequences and variant impact, it is important to choose a method that is appropriate for the data used in prediction. For the purposes of training a tissue-specific model to predict impacts on longer non-coding sequences, approaches such as a gapped k-mer support vector machine (GKM-SVM) can effectively predict the functional impact of single nucleotide variant impacts within CREs (deltaSVM).^28-30^ This GKM-SVM approach has been applied successfully to predict sequence values in the context of specific mouse retinal enhancers.^31-34^ However, to date, it has not been applied across a wider set of human retinal epigenomic data to perform a comprehensive prediction of CRE variant impact scores.

In this study, we applied GKM-SVM modeling with variant impact score prediction (deltaSVM) in a high-throughput manner to predict the functional impact of variants in human retinal CRE sequences. We generated adult human retina ATAC-seq data to determine a high-confidence set of 18.9k putative CREs.^35^ We then used GKM-SVM to train a model which specifically distinguishes retinal CREs versus genomic background sequences, while reserving 20% of candidate CRE sequences as a hold-out dataset for model testing. We then performed *in silico* saturation mutagenesis on this hold-out dataset to generate a database of all possible single nucleotide variants (SNVs) for 3,773 test CREs. We compared these variants to the reference sequence via deltaSVM, generating impact scores for each potential variant. The model revealed that predicted impact scores correlate with allele frequencies in human sequences, and with phylogenetic conservation within candidate CREs. Additionally, we observed distinct negative prediction scores when a variant disrupted the core sequence of a known retinal TF binding motif, consistent with a putative deleterious effect. As a further demonstration of functional relevance, this model was able to predict the consequences of sequence variations when compared to a mutational scan of the mouse Rhodopsin promoter,^36^ showing that the model is robust even across species. Using a larger set of putative retinal CREs, we generated a database of variant impact scores in ocular non-coding sequences (VISIONS) available on the UCSC genome browser. This analysis could be used to identify non-coding variants with higher disease relevance in the retina and prioritize these alleles for functional follow up. By addressing this diagnostic gap, we aim to contribute to a more robust elucidation of CRE function in the human retina.

## Methods

### Input Data Sources

For positive training data, we generated ATAC sequencing datasets from adult human retinas as previously reported (Figure 1B).^35^ These data and other related datasets have been assembled in a searchable track hub on the University of California, Santa Cruz (UCSC) genome browser (https://tinyurl.com/CherryLab-EyeBrowser).^35^ Raw data files were aligned to the hg38 reference genome using Burrows-Wheeler Aligner (BWA), and file format conversions were carried out using SAMtools and BEDtools.^38-40^ Peaks were then called on each dataset using the Model-based Analysis of ChIP-Seq (MACS2) algorithm.^41^ The Irreproducible Discovery Rate (IDR) workflow was implemented to generate a high-confidence set of 18.9k summits of accessible regions by ATAC-seq.^42^ Summits were all extended ±150bp to generate a set of 18,866 putative CRE regions. For the purposes of training and validating the primary model, 80% of peaks were randomly selected for training, and the remaining 20% were used as hold-out data to test the model.

**Figure 1.**
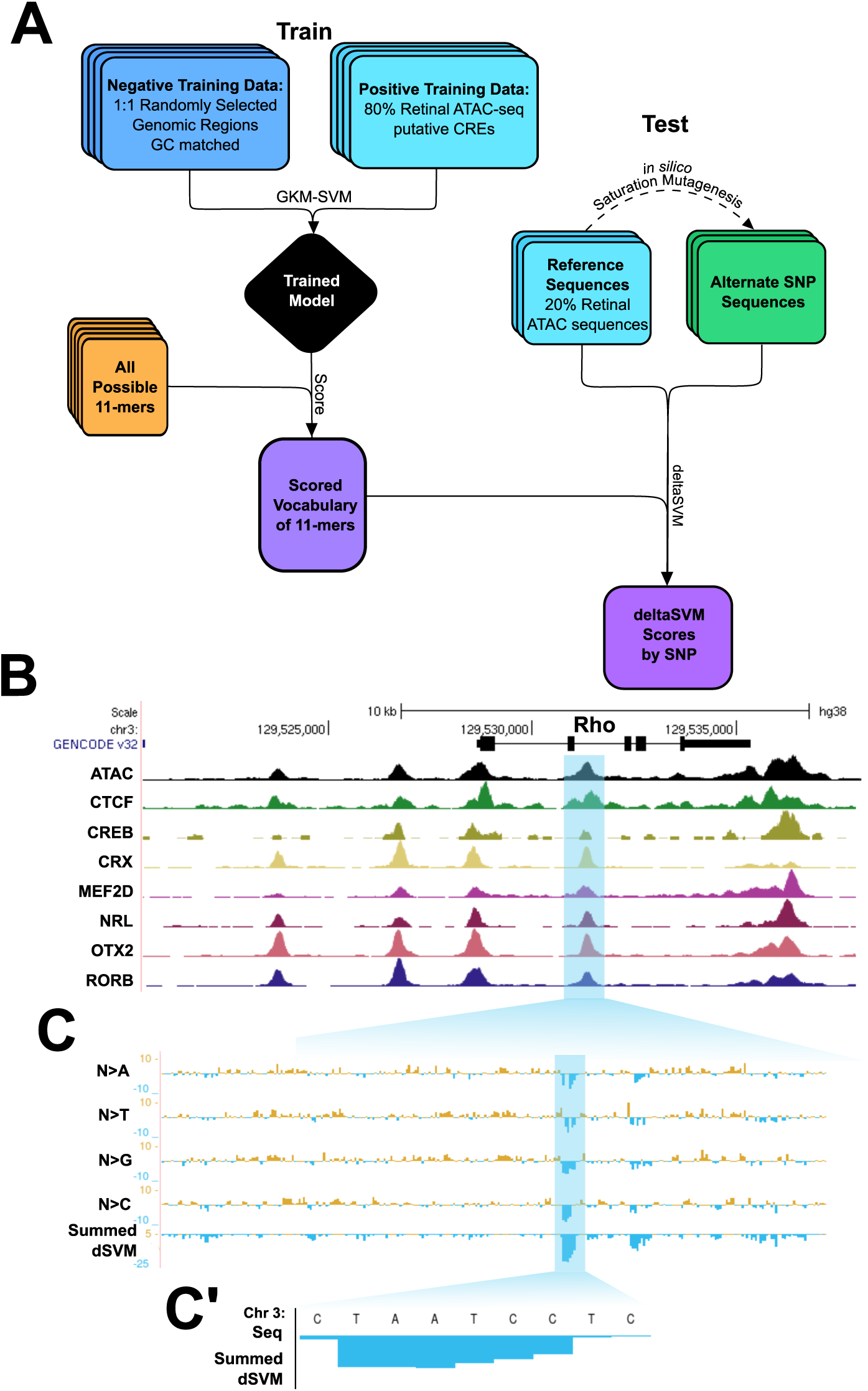
Model Overview and Training Data A.) A schematic overviewing the workflow used in this study in generation of a GKM-SVM based model trained on retinal data and randomly selected genomic regions, and deltaSVM variant impact scores generated through model ranking of *in silico* saturation mutagenesis of putative retinal CREs. B.) UCSC genome browser track positioned at the Rhodopsin (Rho) gene visualizing tracks of the ATAC and ChIP-seq datasets used to generate the Positive Training dataset, schematized in (A). One selected region of interest highlighted in blue (C). Within the highlighted region in (B), base-pair resolution deltaSVM variant impact scores, separated by bp substitution, and summed negative scores. A region of continuous negative scores is highlighted in blue. (C’). In the highlighted region from (C), the summed deltaSVM scores highlighting the core TAATC motif of the OTX2 binding site.

For comparisons to non-retinal data, published data from the GEO database were used including ATAC-seq datasets from Retinal Pigmented Epithelium (RPE),^43^ Primary Visual Cortex (PVC),^44^ and Lung Fibroblasts.^45^ Raw data files were processed as with retinal data, and peaks were called with the same parameters using the MACS2 algorithm and the IDR workflow, and summits were extended ±150bp. For comparisons to retinal data, all regions overlapping with the retinal ATAC peaks were removed using BEDtools intersect. Comparisons of deltaSVM scores to reporter assay expression were made relative to mouse expression data of saturation mutagenesis in RhoCRE3 from Kwasnieski et al.^36^

### SVM Model Training and Validation

To train an SVM model in a biologically meaningful way, the positive training data described above must be compared to an appropriate negative training set. To generate a negative training dataset, 1,000,000 regions were randomly selected from the hg38 genome and extended to 301 bp. These regions were filtered against our positive training data, using BEDtools intersect to eliminate any overlapping sequences (Supplementary Figure 1). From here, an equal number of 301 bp sequences that do not overlap the positive training regions were randomly chosen and GC-matched to the positive set using oPOSSUM.^46^

After selecting the training regions, genomic coordinates were converted to fasta format with BEDtools and used to train a model using LS-GKM gkmtrain, developed by Lee, Beer, and colleagues.^47^ The SVM was trained with the gkmtrain hyperparameters: L=11, k=7, d=3, C=1, t=2, e=0.005; adopted from Shigaki et. al (Figure 1 A).^30^

To validate the classification of the model, the training data was used in a 5-fold cross validation using gkmtrain -x 5 -L 11 -k 7 -d 3 -C 1 -t 2 -e 0.005 to generate performance prediction scores of all regions. Model accuracy was visualized using this data in Receiver Operator Characteristic and Precision-Recall curve graphs as calculated using the ROCR package.^48^ To assess the parameters and results of this primary model, an additional control model was trained with the same parameters and on the same data, but where the positive and negative labels were randomly shuffled to understand the false discovery rate of the GKM-SVM. To compare genomic region performance in the model, retinal and non-retinal genomic peaks were scored using LS-GKM’s gkmpredict. Because GKM-SVM scores in many samples were non-normally distributed, significant differences between retinal and non-retinal data was scored by Kruskal-Wallis chi-squared test, and Pairwise Wilcoxon rank sum test for individual comparisons.

### Vocabulary and Sequence Scoring

To build a regulatory sequence vocabulary, all possible 2,097,152 non-redundant 11bp sequences (11-mers) were generated using *nrkmers*.*py* from LS-GKM and scored by the trained SVM model using *gkmpredict*. To validate the biological relevance of the vocabulary scores, scores were sorted by gkmpredict score value. The top and bottom 1% of scored 11mers were subset for validation and validated against known TF binding motifs.

### Variant Impact Scoring on *in silico* Saturation Mutagenesis

With the previously defined 20% holdout set of 3,773 regions, we scored putative SNVs. To simulate a deep mutational scan, we conducted *in silico* saturation mutagenesis with a custom-made script, yielding 4,542,692 computationally generated sequences that each contained exactly one regulatory SNV. The *deltaSVM*.*pl* script was used to quantitatively assess these variant sequences relative to the consensus allele by referencing the regulatory sequence vocabulary, allowing for the calculation of variant impact scores at a single base resolution.^47^

### deltaSVM Variant Impact Validation

To assess the biological relevance of deltaSVM scores, scores were correlated against human population allele counts. Bcftools^49^ was first used to query the Genome Aggregation Database (gnomAD v3) for indel variants that overlapped with CRE windows defined by the prediction set.^50^ For each 301 bp CRE window, BEDtools was used to compute base-wise summary metrics for indel variant counts and variant impact respectively by summing allele counts per base and negative deltaSVM scores per base. These summary metrics were averaged across CREs to map out the corresponding positional profiles. The relationship between summary metrics was quantified with a Pearson correlation metric between average variant impact scores and indel counts.

### PhyloP Conservation Scores

Phylogenetic P values (PhyloP) of conservation from the PHAST package^51^ across 20 mammalian species were downloaded from the UCSC genome browser. The deltaSVM scores were correlated with PhyloP conservation scores. Alleles were binned into representative groups of 2000 alleles from the top, middle, and bottom-most deltaSVM scores for plotting of conservation. Because PhyloP scores in deltaSVM bins were non-normally distributed, significant differences between bins was scored by Kruskal-Wallis chi-squared test, and Pairwise Wilcoxon rank sum test for individual comparisons.

### Transcription Factor Motif Analysis

Positive training data was scored for TF motif enrichment using HOMER findMotifsGenome.pl and findMotifs.pl.^52^ Common retinal motifs were selected from the known motif results for analyses of model relevancy.

For the scoring of motif prevalence in distinct sequences rather than overall enrichment in a set, sequences were scored against the Homo Sapiens Comprehensive Model Collection (HOCOMOCO) v11 Core database.^53^ Motif prevalence in vocabulary and deltaSVM bins against the HOCOMOCO v11 database were scored using Find Individual Motif Occurrences (FIMO) from the MEME suite of tools with a significance threshold of *P* ≤ 1 *x* 10^−4^.^54^ Significant changes in average dSVM within Motifs by bp was scored by ANOVA with post-hoc Tukey test.

For the validation of motif interference in deltaSVM scores, known motif positions were obtained from HOMER and positions were extended by 25bps using Bedtools slop. Bedtools intersect was used to identify motifs that were with regions of interest and collect corresponding average delta-SVM scores.

## Results

### Human Retinal Epigenomic Data Can Be Used to Train a GKM-SVM Model

To generate impact score predictions for single nucleotide variants within human retinal CREs, we first trained a gapped k-mer support vector machine model (GKM-SVM)^28^ to evaluate putative CRE sequences (Figure 1A). As input, we started with a set of genomic windows defined by high confidence ATAC-seq DNA-accessibility peaks. We split this set such that 80% of ATAC regions (∼15k candidate CREs) were used as a positive training set, and 20% (∼3.7k) were kept as a hold-out set to test the validity of predicted impact scores. (Figure 1B, Supplementary Figure 1). Also for input, we generated an equal sized negative training set of GC-matched non-coding genomic sequences (Supplementary Figure 1B & C). As expected for putative CREs and control regions, we found that both positive and negative datasets were enriched for intronic and intergenic regions. We also found that the negative training dataset was depleted of promoter regions when we removed any overlap with the positive training data. As a control, we trained a separate model using the same input data but with the positive and negative region labels shuffled randomly, to demonstrate the baseline behavior of the model parameters.

To use our trained model to predict the impact of CRE variants, we next generated a scored vocabulary of all possible non-redundant 11mer sequences. This k-mer length was chosen because it is long enough to encompass most eukaryotic TF binding motifs.^55^ We then used the trained model to weigh each 11-mer based on its relative similarity to the positive training set (positive values) versus the negative training set (negative values). This scored vocabulary was subsequently used to evaluate variant sequences in the generation of variant impact (deltaSVM) scores (Figure 1A).

Finally, to generate individual CRE variant impact scores, we performed base-wise *in silico* saturation mutagenesis on CREs from the 20% hold-out dataset. We then used the scored 11-mer vocabulary to predict impact scores for every possible single nucleotide variant within these CREs. These predicted impact scores represent the difference between the sum of all 11-mers that scan across a given single nucleotide variant compared to the sum of those that scan across the reference allele (Figure 1A, 1C).^29^ A negative impact score therefore is assigned to a variant when it causes the sequence to become less similar to the positive training dataset compared to the original reference sequence. When deltaSVM scores are combined across a genomic region, distinct features of CREs become apparent. For example, inspecting for contiguous, highly negative summed deltaSVM scores, it is possible to identify well-characterized TF binding motifs in the reference sequence, such as the TAATCC motif favored by the K50 homeodomain transcription factors OTX2 and CRX and the CTCF binding motif (Figure 1C, Supplementary Figure 3).

### Performance and Biological Relevance of the Trained SVM Model

To assess the validity of this approach, we first performed 5-fold cross validation on the original and shuffled models. The training data was randomly assigned to one of five outgroups, and each outgroup was scored against a model trained excluding that outgroup. This cross validation allows for the specific calculation of false positives and negatives, as well as model precision. These measures of model accuracy can be plotted as a Receiver Operating Characteristic (ROC) curve (Figure 2A), or a Precision-Recall Curve (Figure 2B). For this model’s ROC curve, an area under the curve (AUC) of 0.951 was achieved, indicating a highly accurate model with low false positivity. Similarly, the precision-recall curve for this model demonstrated an AUC = 0.956, indicating high precision. In contrast, when positive and negative labels were shuffled for the training data, the ROC and precision-recall curves demonstrated baseline AUCs, showcasing the specificity of model training gained by the true positive and negative datasets.

**Figure 2.**
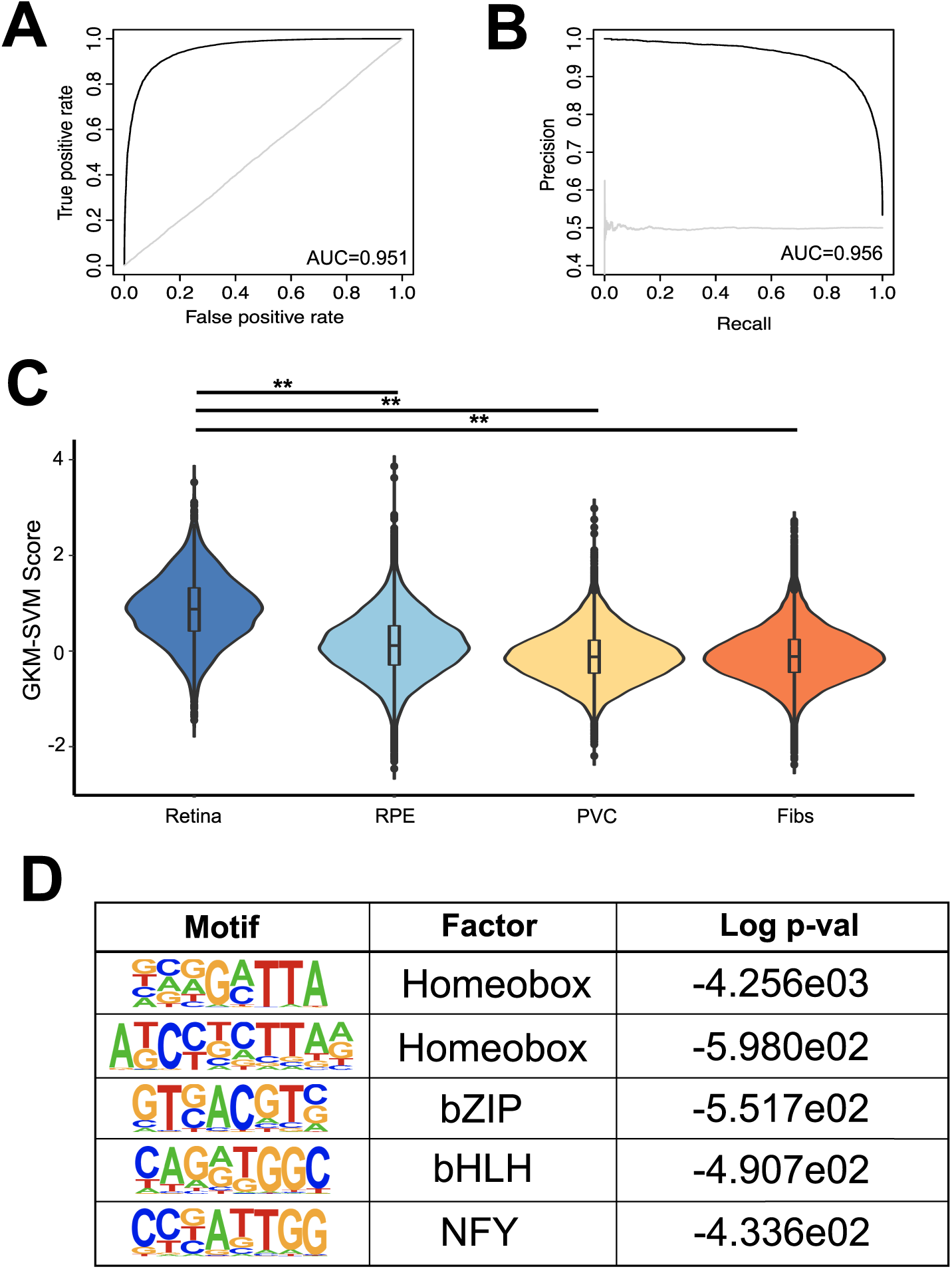
The GKM-SVM model is Accurate, and Retinal-Specific A. Receiver Operating Characteristic curve for 5-fold cross-validation of the GKM-SVM model trained on retinal epigenomic data (black) and for the model trained on shuffled positive and negative training data (light grey). Area under the curve (AUC) ATAC = 0.951; shuffled = 0.498. Precision-Recall curve for 5-fold cross-validation of the GKM-SVM model trained on retinal epigenomic data. AUC ATAC = 0.956, shuffled = 0.499. C. Violin plot demonstrating GKM-SVM model scores for Retinal positive training data, Retinal Pigmented Epithelium ATAC-seq peaks,^41^ Primary Visual Cortex ATAC-seq peaks,^42^ and human Fibroblast ATAC-seq peaks.^43^ Kruskal-Wallace p < 2e-16. For all pairwise comparisons to Retinal ATAC by Wilcoxon rank sum, p < 2e-16 (**) Bonferroni adjusted. D. Top enriched TF motifs from HOMER in top 1% of scored model 11mer vocabulary,

To determine the tissue-specificity of our trained models, we next used these models to compare retinal versus non-retinal CRE sequences. We found that our original model scored retinal-specific CREs from our hold-out dataset much more highly than non-retinal CRE datasets of equal size (Figure 2C). Retinal ATAC-seq regions demonstrated a wide variety of scores, averaging at a GKM-SVM score of 0.870. This was significantly higher than all other non-retinal ATAC-seq data (p values < 2e-16 in pairwise Wilcoxon rank sum tests, Bonferroni adjusted). Retinal pigmented epithelium (RPE), being developmentally related to the retina, scored most neutrally with an average score of 0.125, as compared to the primary visual cortex (PVC) at -0.104, and fibroblasts at -0.087 (Figure 2C). These differences were eliminated when we used the shuffled model to score CREs, demonstrating the specificity of our original model for evaluating retinal CREs.

To further assess the tissue-specific relevance of our original model, we searched for the enrichment of known transcription factor binding motifs within the top 1% of the scored 11-mer vocabulary. Within this group we found significant enrichment for motifs shared by well-known retinal transcription factors (Figure 2D). Photoreceptor-associated motifs such as homeobox domain motifs consistent with CRX and OTX2 binding were most highly enriched, while more broadly expressed retinal TF motifs such as bHLH motifs were also highly ranked. This enrichment within the 11mer vocabulary suggests an additional level of tissue-specificity in our model.

### Variant Impact Scores Correlate with Conservation of Non-Coding Sequences

Pathological variants within human retinal CREs are relatively rare but can disrupt visual function.^2,12-14^ We therefore reasoned that if our variant impact scores were biologically relevant, then strongly scored variants should be rare within the normal human population. To test this, we compared our predicted variant impact (deltaSVM) scores to allele frequency in the Genome Aggregation Database (GnomAD).^50^ When directly plotted against each other, we found that alleles with high frequencies in the population clearly aggregated around the neutral deltaSVM scores (Figure 3A, A’). Conversely, alleles with a large predicted impact (negative or positive) are not frequently found in the population. The majority of alleles scored with a deltaSVM between -3 and 3, with 1% of all alleles scoring less than -4.9, and another 1% scoring more than 3.4 (Figure 3B). By comparison, when constrained to deltaSVM scores from -2.5 to 2.5 to select more common alleles, 1% score bins span respectively from -2.5 to -2.3, and 2.26 to 2.5. These four 1% bins are defined as the top and bottom 1% and the mid -top and -bottom 1%, as shown in Figure 3 A and B. A fifth bin was additionally defined, spanning the most neutral deltaSVM scores from -0.001 to 0.001 (Figure 3A, A’). This comparison suggests that variants predicted to have a strong impact on CRE function may be deleterious because they are rare in the human population.

**Figure 3.**
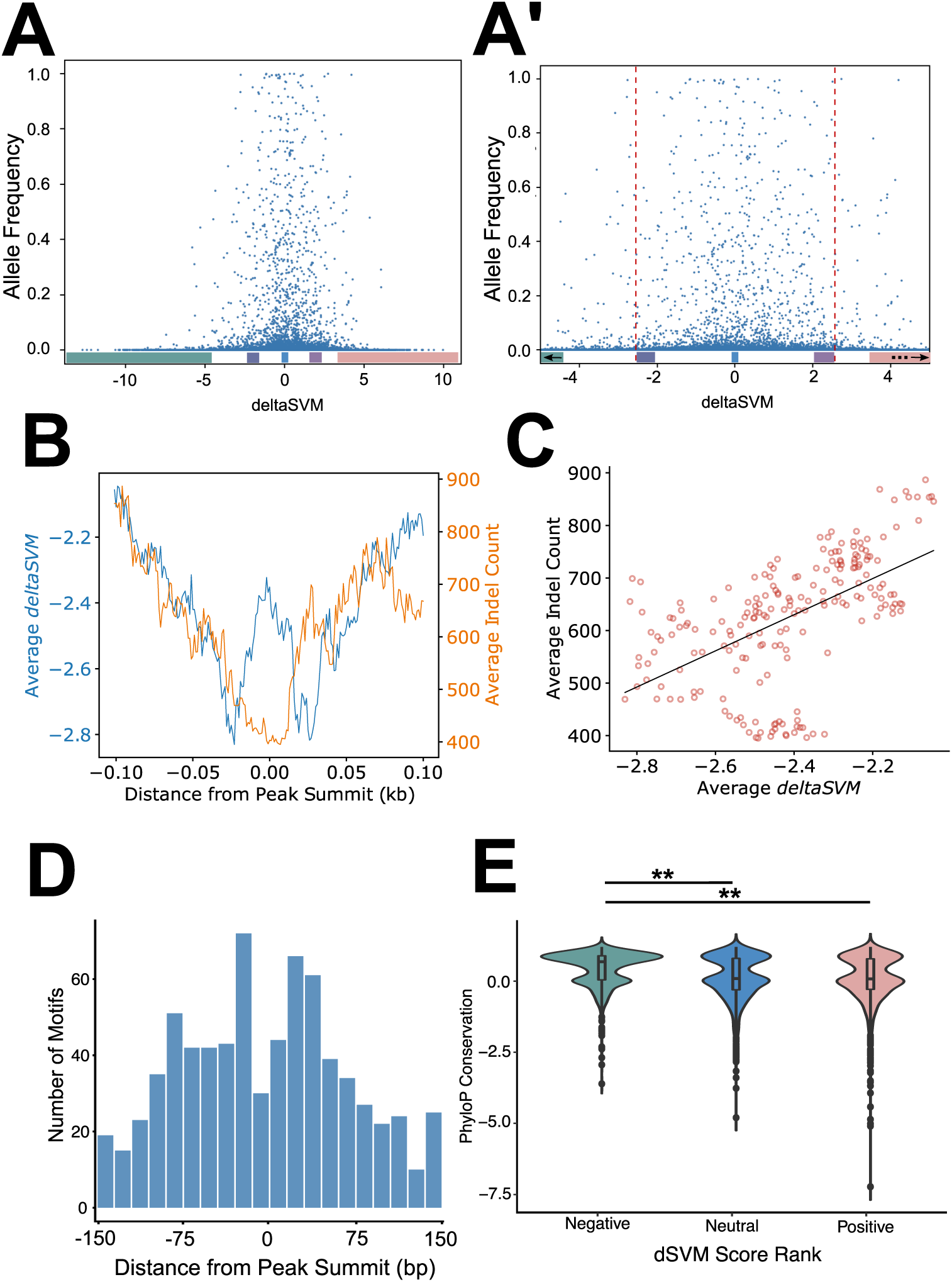
deltaSVM Scores Match Allele Frequencies, Conservation A. Scatterplot demonstrating the correlation between deltaSVM scores and allele frequencies from the GnomAD database. deltaSVM bins for the bottom, mid-bottom, mid-top and top 1% of scores are highlighted along the X axis. A’. Alternate view of (A), demonstrating values from deltaSVM scores -4 to 4. deltaSVM bins for the bottom (partial), mid-bottom, neutral, mid-top and top (partial) 1% of scores are highlighted along the X axis. Arrows indicate points beyond the axis limits for the highlighted bins. B. Changes in average deltaSVM and Indel counts across the average 301 bp window of 20% outgroup CREs. C. Correlation between average deltaSVM and Indel count by position along the averaged 301 bp window. Pearson correlation 0.303 D. Sums of retinal motif classes across the 301 bp window of 20% outgroup CREs in 15 bp bins. E. Violin plot of PhyloP conservation scores in Negative, Neutral, and Positive deltaSVM scores (Kruskal-Wallace p < 2e-16, Wilcoxon rank sum: **: p < 0.0001, Bonferroni adjusted).

Another test of the relevance of the variant scores is the distribution of these scores across the linear sequence of CREs. The center of retinal CREs is typically depleted of indels in the normal human population where indels could disrupt transcription factor binding or the spacing between motifs.^35^ We would expect a similar trend for our predicted impact scores where deltaSVM values would be more strongly negative toward the center of a CRE. We therefore averaged negative deltaSVMs across all 3.7k CRE windows and compared them to indel count and position within CREs. When plotted, a clear trend emerges, where deltaSVM scores, as well as the average indel count, decreased toward the center of CREs, corresponding to the peak of DNA accessibility (Figure 3B). These deltaSVM scores correlated with the frequency of indels (Pearson Correlation Coefficient = 0.685, Figure 3C). However, at the center of the CRE, the average deltaSVM score increased locally, generating a bimodal distribution of negative impact scores. This may reflect the actual distribution of TF binding within CREs. To test this, we analyzed the distribution of TF motifs and found a similar trend. Motifs consistent with TFs such as CREB, CRX, MEF2D, NRL, OTX2, and RORB were most enriched directly adjacent the center of CREs (Figure 3D, Supplementary Figure 4). Together these results suggest that variants near, but not at, the DNA-accessibility-defined center of CREs are likely to have a negative impact on CRE function, especially in regions of TF binding.

Consistent with these trends, we would expect deltaSVM to be negatively correlated with conservation values across species. Evolutionary conservation of specific CRE sequences suggests that those sequences are functionally important. To test this, we binned deltaSVM scores into the most negative, most positive, and neutral deltaSVM categories as defined in Figure 3A, and compared these categories to phylogenetic conservation scores from the PhyloP database. The most negatively scored SNVs (average -9.98) corresponded to more conserved sequences (higher PhyloP scores around an average of 0.456), while more neutral (average 4.9e-06) or positively (7.39) scored SNVs had lower conservation scores (Neutral average PhyloP score = 0.119, Positive average PhyloP score = 0.101, Negative to Neutral/Positive p values < 2e-16, Bonferroni corrected, Figure 3E). SNVs with more negative predicted impacts therefore appear to be more highly conserved, indicating their potential regulatory value in a given putative CRE.

### Highly Negative Variant Impact Scores Disrupt TF Binding Motifs

The correspondence of variant impact scores with allele frequency and conservation suggests that deltaSVM value correlates with TF binding motifs. To evaluate this directly, we first determined the counts of specific TF binding motifs in the most negative, most positive, and the neutral deltaSVM categories. In each of these categories (Figure 3A), the 22 bp around a given SNV were scored for the presence of known motifs in the HOCOMOCO human motif database.^53^ In the most negatively scored category, we found many well-characterized retinal TF motifs, such as OTX2 and CREB represented in the reference sequences (Figure 4A, A’’, Supplementary Figure 5). By contrast, in the SNV sequences there were far fewer motifs as scored by FIMO, indicating that the variant sequences specifically disrupt the sequence of the motif. This pattern varies in the mid-bottom deltaSVM scoring variants. While MEF2D motifs show a high number of motif calls, OTX2 calls by contrast are much less frequent (Figure 4A’). The variants scored in the top bins demonstrate the opposite trend, with few calls for motifs of interest in the reference sequence, with modest increases in the variants likely due to situations where the variant coverts a sequence into an approximation of a TF binding motif (Figure 4A, A’, A’’, Supplementary Figure 5).

**Figure 4.**
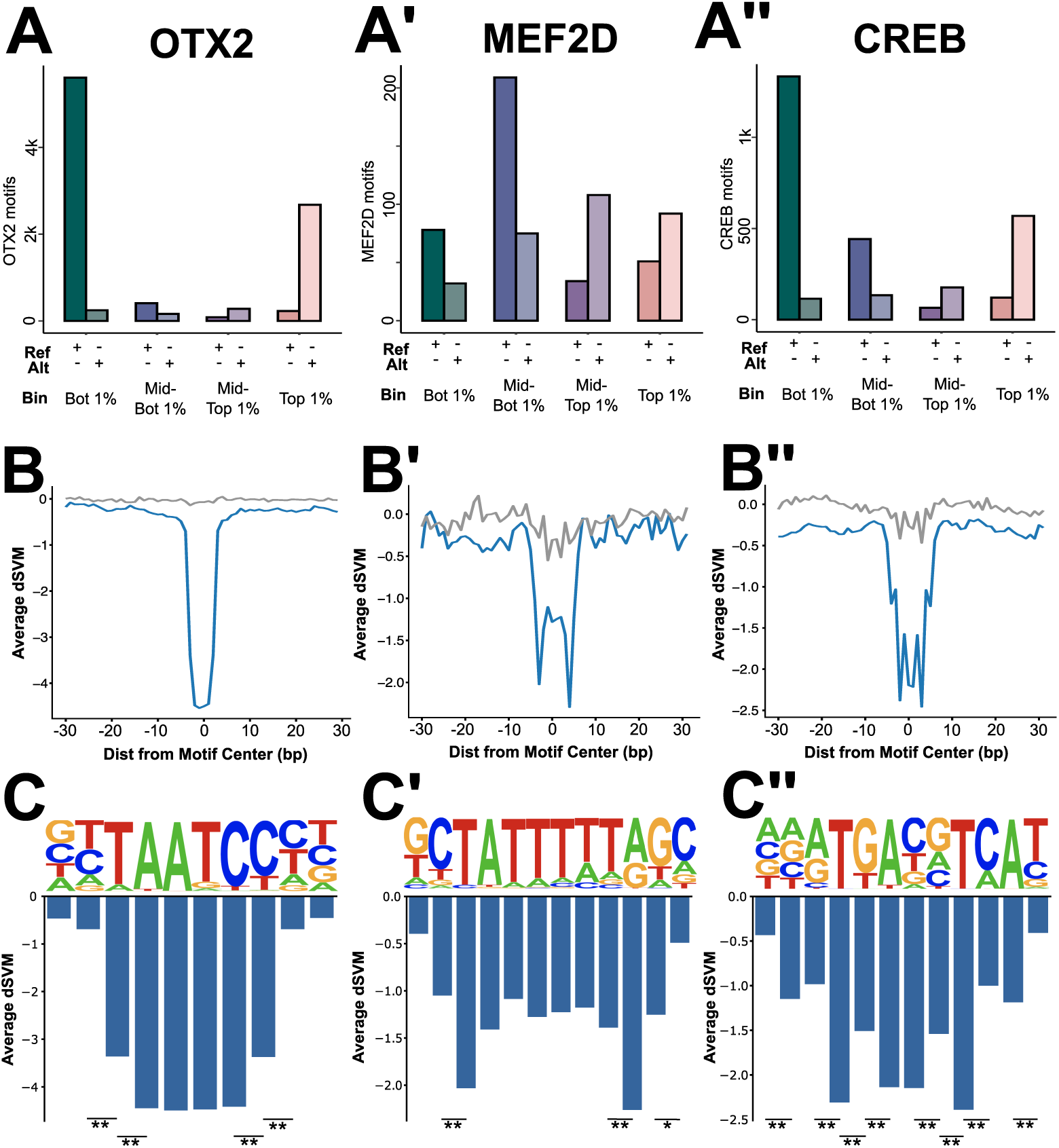
Disruption of Retinal TF Motifs Dramatically Reduces deltaSVM Score A-A’’. Numbers of motifs as scored by FIMO in reference and variant sequences for bins highlighted in (3A). Motifs shown are OTX2 (A), MEF2D (A’), and CREB (A’’). B-B’’. Line plots showing the average deltaSVM for SNVs +/- 25 bp around the core motifs shown in (A-A’’) in blue. Scores for the same motifs in the shuffled model in grey. C-C’’. Bar plots showing the average deltaSVM for SNVs on a base pair resolution within the core motifs of those shown in (A-A’’). (ANOVA with Post-hoc Tukey: *: p<0.02; **: p< 0.002)

To gain a better understanding of the relationship between TF motifs within CREs and our predicted variant impact scores, we identified all instances of specific TF motifs in our test data set and centered these on 60bp windows. We then plotted the distribution of deltaSVM scores across these windows. We observed that the scores dip dramatically around canonical motif sequences while the flanking regions are relatively unaffected (Figure 4B, B’, B’’, Supplementary Figure 6). This indicated that our scoring strategy is uniquely sensitive to these motifs. TF motif sequences however allow for flexibility at specific positions across the motif. We therefore sought to determine how impact scores varied within a motif itself. At a single base pair resolution, we found that the significance of the core motif of some TFs such as OTX2 is apparent (Figure 4C, Supplementary Figure 6. Average deltaSVM scores for SNVs in the core TAATCC sequence are more negative than for SNVs in contextual positions immediately adjacent (Figure 4C). For other motifs, changes to key nucleotides in a motif sequence become apparent, with larger decreases highlighting the CTA/TAR caps of the MEF2D consensus motif (Figure 4C’) as well as key bases in the CREB motif (Figure 4C’’). These changes in deltaSVM scores along TF binding motifs demonstrate the specificity of these scores to isolate crucial core sequences in a putative CRE, and where alterations to the sequence may have significant impact on function.

### Prediction Scores Across a Conserved CRE Match Changes in Reporter Expression

Prior studies have demonstrated the ability to experimentally test the impact of every possible SNV within a retinal CRE using a massively parallel reporter (MPRA)-based approach.^36^ Kwasnieski et al. used SNV saturation mutagenesis of the mouse Rhodopsin promoter to test the impact of every possible variant with base-pair resolution (Figure 5A).^36^ This analysis highlighted the unique importance of CRX and NRL motif sequences within the larger CRE. As a final test of our predicted variant impact scores, we used our human retinal CRE-trained model to assign predicted variant impact scores every possible SNV within this mouse sequence (Figure 5B). Although the human ortholog of this CRE was not included in the original 80% training set and despite being tested against reporter data generated in the mouse retina, the model predicted markedly negative deltaSVM scores overlapping the previously identified TF binding sites, highlighted in figure 5A and 5B, as well as similarities in the region between the CRX(2) and NRL binding motifs (Figure 5B). When relative expression from 5A was plotted against deltaSVM scores in 5B, these values were found to be positively correlated, with a Pearson correlation coefficient of 0.506. Altogether, this correlation and consistency across TF motifs, suggested to us that our CRE variant prediction strategy is robust.

**Figure 5.**
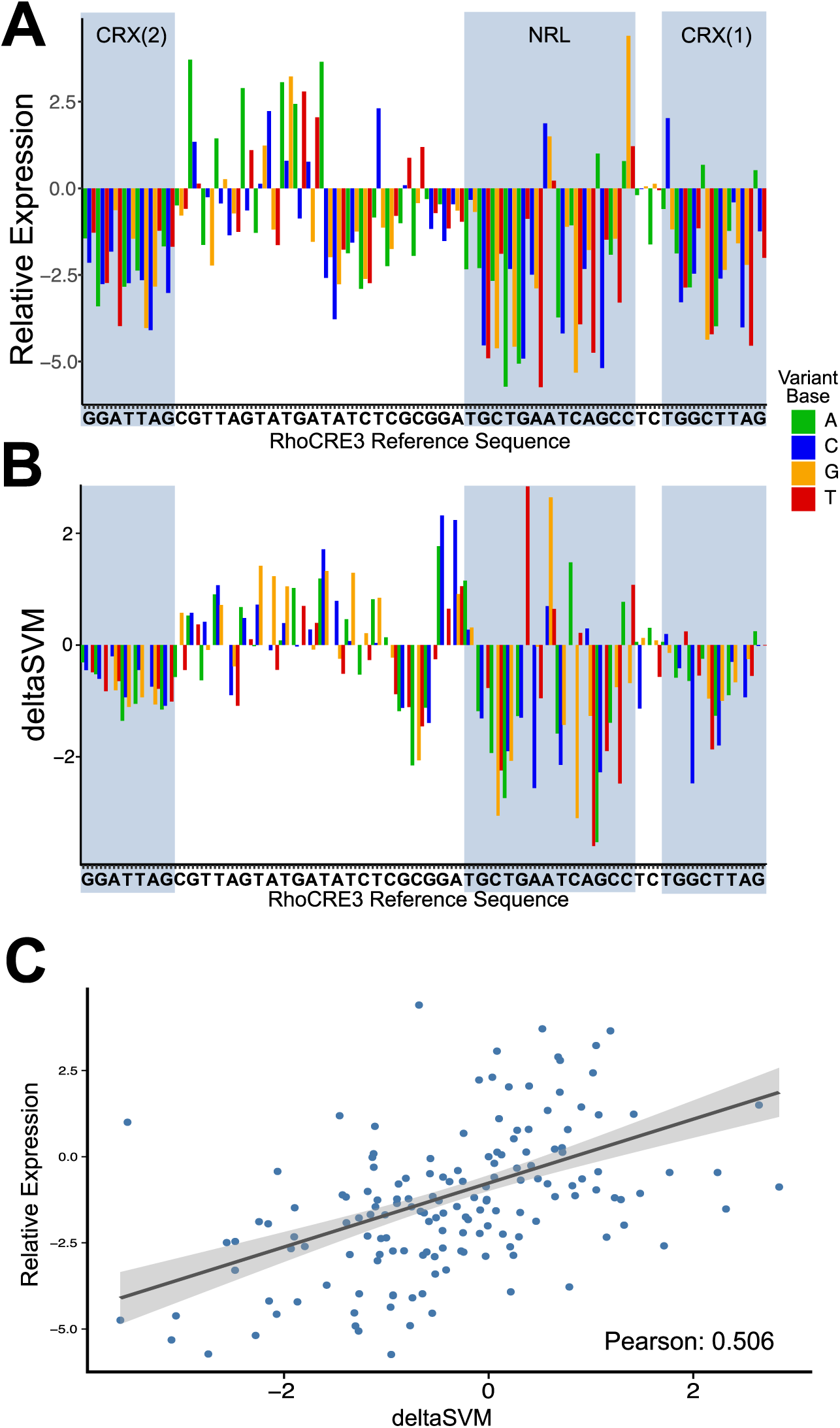
deltaSVM Scores for SNVs in RhoCRE3 Correlate to Changes in Reporter Expression. A. Relative expression (log2(mut/wt)) of fluorescent reporter for mutations in RhoCRE3 in mouse retina from Kwasnieski et al.^34^ Identified TF binding sites of CRX (1 and 2) and NRL are highlighted. B. deltaSVM variant impact scores of the same SNVs as in (A) along the RhoCRE3 locus. Identified TF binding sites of CRX (1 and 2) and NRL are highlighted. C. Scatter plot of relative expression and deltaSVM scores in (A) and (B) with linear regression and 95% confidence intervals. Pearson = 0.506.

### A Resource for Human Retinal Regulatory Variant Interpretation

The analyses described above suggested that this variant impact scoring strategy using the GKM-SVM/deltaSVM workflow trained on human retinal ATAC-seq data has several biologically relevant features. We therefore extended these scores to include a more inclusive set of ∼39k putative retinal CREs as determined by our adult human retinal ATAC-Seq analysis. This analysis entitled “Variant Impact Scores in Ocular Non-coding Sequences” (VISIONS) is available on the UCSC genome browser to query and to compare with human retinal DNA-accessibility, transcription factor binding and histone modifications (http://genome.ucsc.edu/s/CherryLab/VISIONS_TrackHub). It is our hope that these predicted impact scores can assist other researchers in identifying and interpreting variants of interest within non-coding retinal regulatory elements.

## Discussion

The identification and characterization of non-coding mutations in retinal *cis*-regulatory elements (CREs) can be a resource-intensive process. This study demonstrates the value of machine learning to identify highly impactful SNVs and to generate an exhaustive analysis of retinal CRE variant impact scores. These scores were generated through training a GKM-SVM model on adult human retinal ATAC data. By utilizing this machine learning-based approach, large epigenomic sequencing datasets can be analyzed, and with the GKM-SVM and deltaSVM approaches, sequence variations can be easily screened. This SVM-based method has been previously used to highlight specific sequence features in the mouse retinal epigenome and to predict retinal-reporter expression post-hoc.^31-34^ Together these previous studies and our current work demonstrate the potential of this approach to characterize human retinal CRE sequences for the identification of crucial features and their variants. While this approach can be applied to many types of sequencing data, the use of general chromatin accessibility via ATAC-seq allows the model to incorporate the sequence features of diverse regulatory elements in a less biased approach than using more specific ChIP-seq data. Ultimately, we hope that the model generated in this study can be used to identify non-coding sequence variants that are likely to disrupt retinal CRE function to guide deeper analyses of non-coding mutations. Currently, single nucleotide variant scores for sequences in 39k putative retinal CREs can be accessed via our UCSC genome browser track to identify variants with large predicted impacts to retinal CRE function (http://genome.ucsc.edu/s/CherryLab/VISIONS_TrackHub).

This study presents a machine learning model of putative human retinal CREs, and the predicted impact of all possible SNVs in a set of tested sequences. This model behaves in a tissue-specific manner, and accurately identifies the enrichment of well characterized TF binding motifs. Through the analysis of related datasets and known motif databases, the model trained on human retinal ATAC data versus genomic background can clearly identify sequences of interest in a biologically relevant manner, specifically scoring retina-associated sequences above non-retinal CRE sequences. Further, in the generation of variant impact deltaSVM scores, the model’s scores follow known conservation, and specifically identify where disrupted sequences intersect canonical transcription factor binding motifs to potentially affect CRE activity. The enrichment of negative deltaSVM scores around known motifs specifically highlights well-characterized core sequences and key base positions in motifs, and thus the ability of the model to recognize the value of these sequences. Additionally, it becomes apparent that distinct motifs contribute differently to model relevancy. Potentially, the motif disruption and severity of the deltaSVM score may be an indicator of the severity of impact on CRE and therefore retinal function. Those SNVs with the most negative deltaSVM scores were associated with the highest level of conservation, demonstrating that these sequences may have a distinct role in retinal function.

When observing these deltaSVM scores in the general context of the CRE, trends become apparent as to where the most impactful variants are found confirming known features of CREs. The trend of deltaSVM scores across putative CREs demonstrates both the known density of true TF binding sites near the summit and the depletion at the summit itself. This is consistent with findings from other studies, which show that TF motifs are most enriched around the center of CREs, but are somewhat depleted at the direct summit.^55^ These data indicate that disruptions to these TF motifs have the most dramatic impact on CRE scoring. Previous studies have performed massively parallel reporter assays to test the function of specific CRE sequences.^36,57,58^ The results of these studies emphasize the impact of specific TF motif disruption and also serve as an important resource for the validation of ML predictions of variant impact. The characterization of these motifs is highly conserved as negative deltaSVM scores from this model specifically correlate with MPRA-based approaches.^30,36^ These results demonstrate both the ability of this model trained on human retinal epigenomic data to identify variants with notable relevance to changes in gene expression, as well as its ability to operate across species in a conserved manner. This ability of the model to identify sequences of interest, especially variants that correlate to losses in gene expression, demonstrates the ability of this model to predict non-coding variants with relevance to retinal disease.

This model has unique value in the retina, in that it can specifically evaluate sequences associated with *cis*-regulatory elements, lending itself to a wide variety of applications. deltaSVM impact scores can be used in the identification of crucial TF binding sites in a high-throughput manner. In particular, the *in silico* saturation mutagenesis approach to generating a database of deltaSVM scores means that variants can be pre-screened by their predicted change in regulatory function. In the screening of regions identified via GWAS, such data can specifically narrow down regions of interest and locations of mutations of functional value to the retina. In more precise applications, variants identified in patients can be quickly ranked by their relevance to this model and prioritized for further functional investigation. This model can be further refined via integration of new epigenomic datasets, in particular single-cell epigenomic datasets to refine the sensitivity and specificity of these predictions. In the rapidly moving field of AI, new machine learning strategies will also likely enable characterization of new and different sequence-based features within CREs.

In sum, this workflow and the resulting prediction scores serve as a promising genomic tool for guiding the interpretation of non-coding sequence variation, and for narrowing the search space for potentially pathogenic regulatory variants in visual disorders. Validation of the model demonstrates its capacity for tissue specificity, and the identification of crucial CRE features. By applying a deltaSVM approach to putative CRE sequences, it is possible to pre-screen variant sequences of interest for further *in vivo* analyses. With further model validation, the presented database of SNV scores could be used in the identification of clinically relevant sequence variations and have applications beyond the bench.

## Acknowledgements

This work was supported by grants from the NIH National Eye Institute R01EY028584 to T.J.C., and the Catalytic Collaboration Grant from the Brotman Baty Institute awarded to T.J.C. and A.Y.L. The authors thank Thomas Vierbuchen, Marty Yang, Eric Thomas, Brendan McShane, and all members of the Cherry Lab for their review and valuable discussions regarding this manuscript.

## Author Contributions

L.S.V., K.L., A.Y.L., and T.J.C. conceived of all experiments and analyses. K.L. and L.S.V. developed ML training and analysis pipeline. L.S.V. performed all final model training. L.S.V. and A.E.T. performed all analyses. Y.W. assisted in data analysis development. S.C. developed UCSC browser track.

## Figures

**Supplementary Figure 1.**
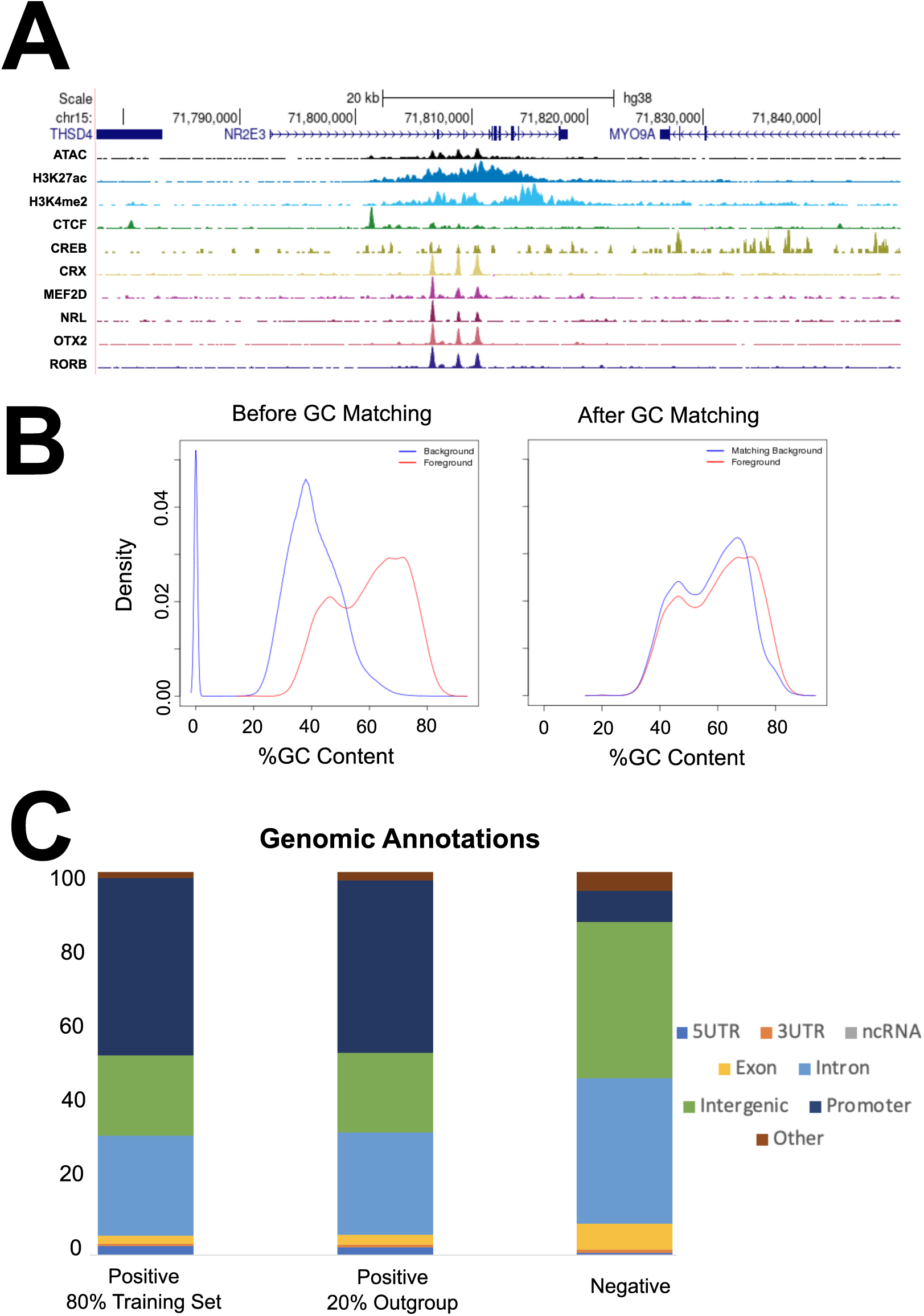
A. UCSC genome browser track positioned at the Nr2e3 gene visualizing tracks of the ATAC and ChIP-seq datasets used to generate the Positive Training dataset, schematized in 1A. B. GC content density of Positive training dataset (ATAC/Foreground in red), and Negative training data (Background in Blue) before and after GC matching with oPOSSUM. C. HOMER genome annotation of the positive (80% training and 20% hold-out) and negative training datasets used to train the GKM-SVM model.

**Supplementary figure 2.**
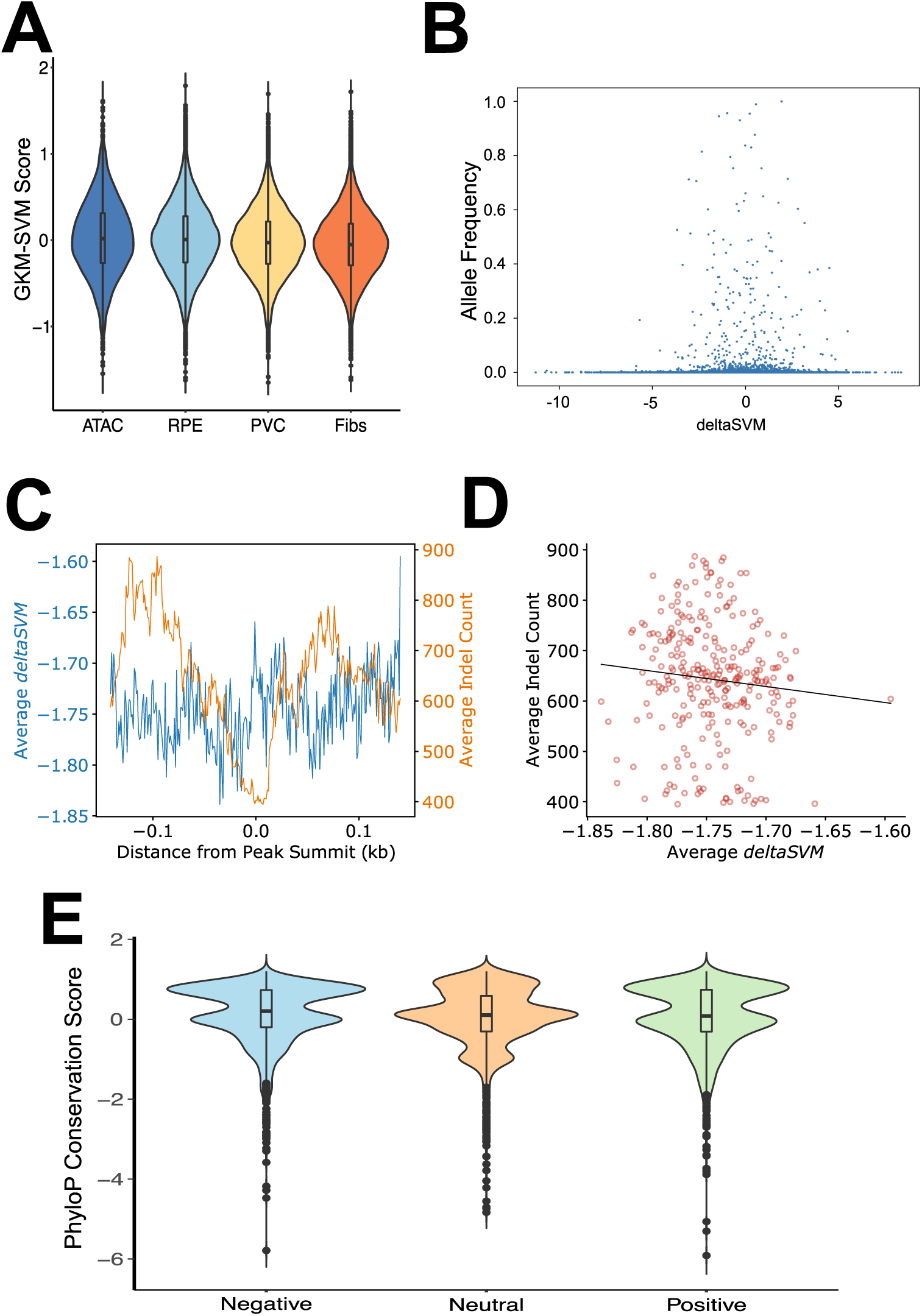
Demonstration of the Shuffled model utilizing ATAC-seq positive and negative training data shuffled randomly. A. Retinal (20% outgroup) and nonretinal sequences scored by the shuffled model. B. deltaSVM scores (in 20% outgroup sequences) for the shuffled model as compared to GnomAD allele frequencies in the same regions. C-D. Average deltaSVM and GnomAD indel counts across the average 301mer in the 20% outgroup CREs demonstrating the trend in score across the CRE (C), and the correlation between average deltaSVM and average indel count (D). E. Violin plot of PhyloP conservation scores in Negative, Neutral, and Positive deltaSVM scores.

**Supplementary Figure 3.**
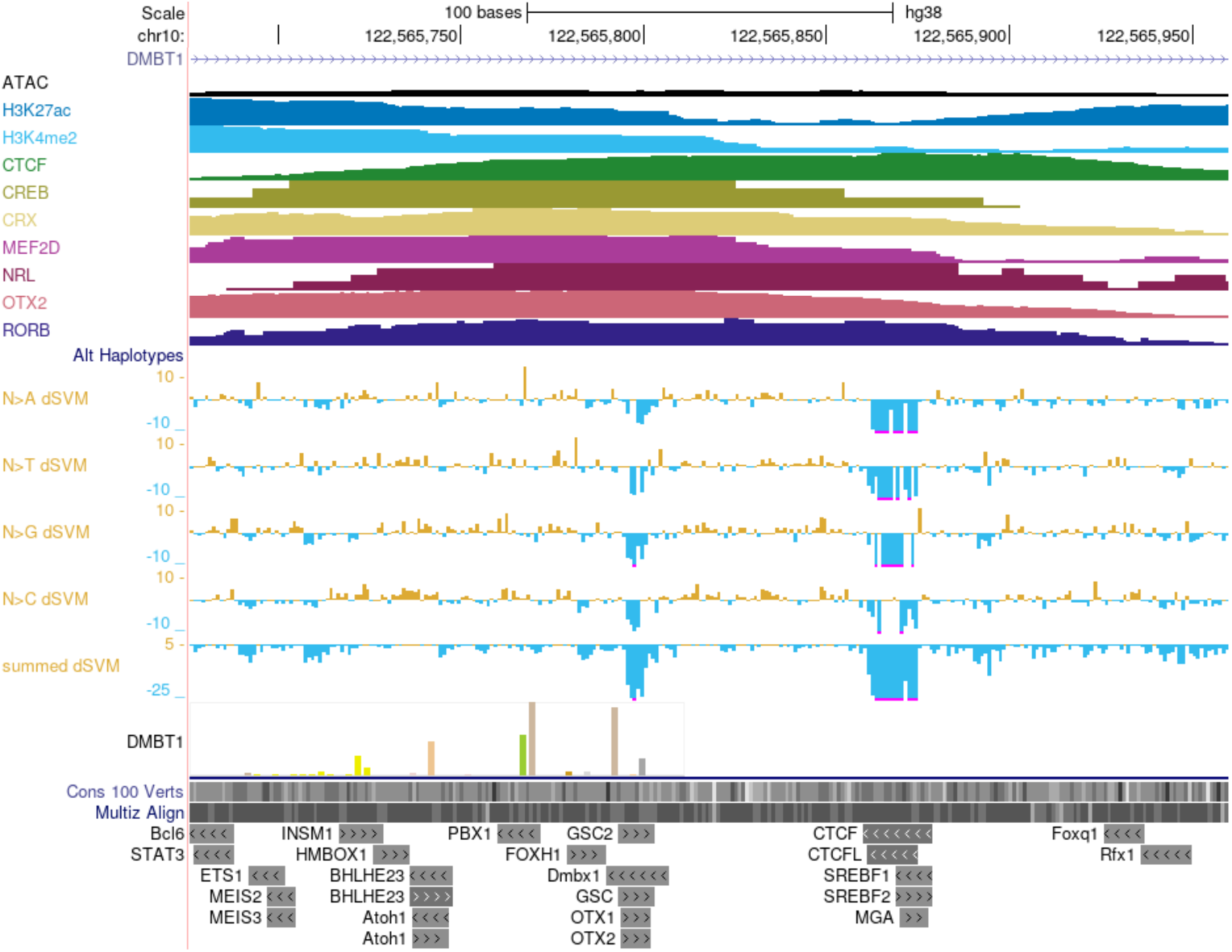
A. UCSC genome browser track positioned at an intergenic putative CRE within the DMBT1 gene visualizing tracks of the ATAC and ChIP-seq datasets used to generate the Positive Training dataset, schematized in 1A, as well as deltaSVM scores split by alternate base substitution and predicted TF binding sites.

**Supplementary Figure 4.**
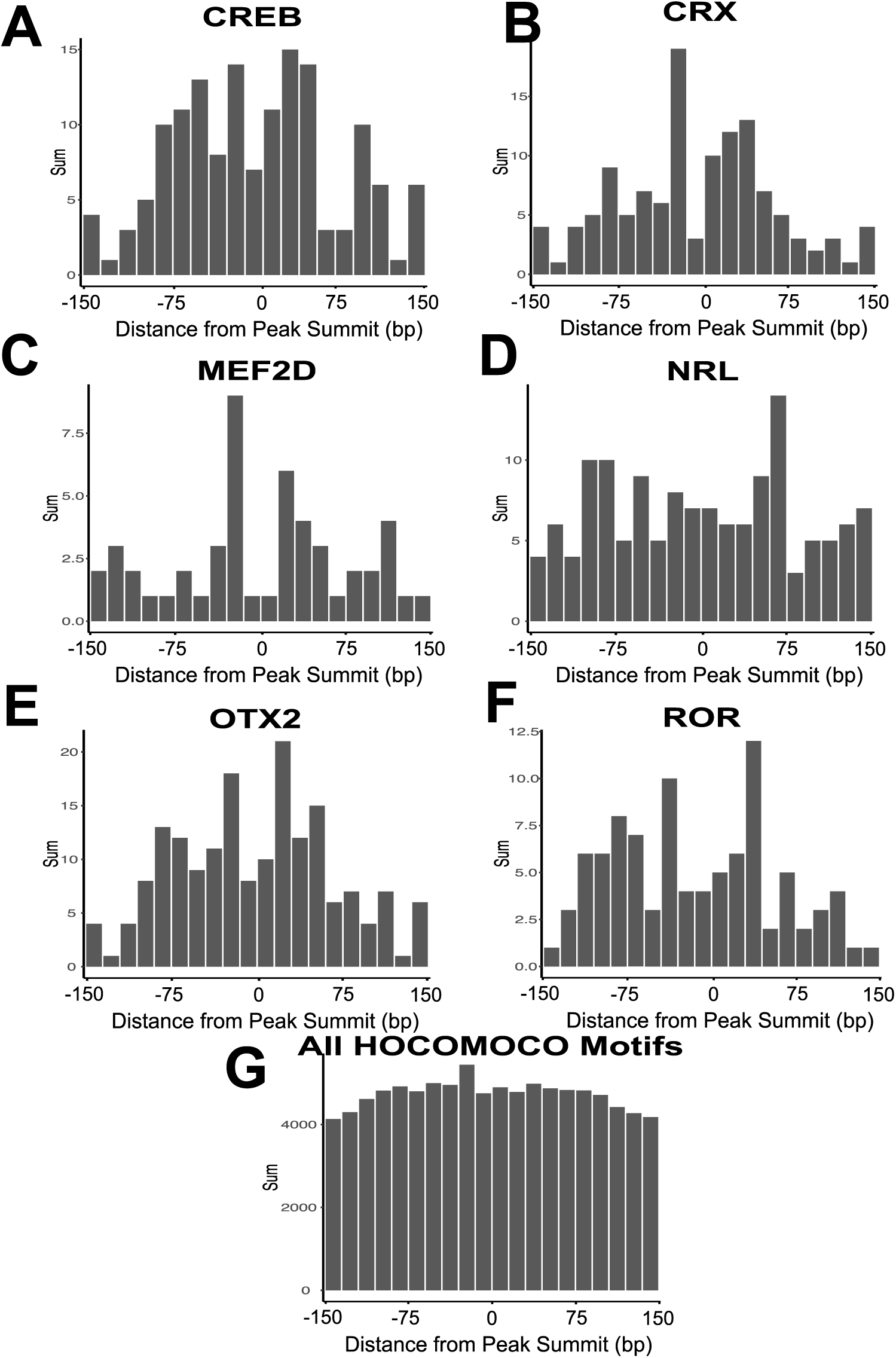
Sums of retinal TF motifs across the 301 bp window of 20% outgroup CREs in 15 bp bins. A-F demonstrate motif sums for individual retinal motifs CREB (A), CRX (B), MEF2D (C), NRL (D), OTX (E), and ROR (F). G. Sums for all possible motifs (retinal and non-retinal) from the HOCOMOCO v11 motif database.

**Supplementary Figure 5.**
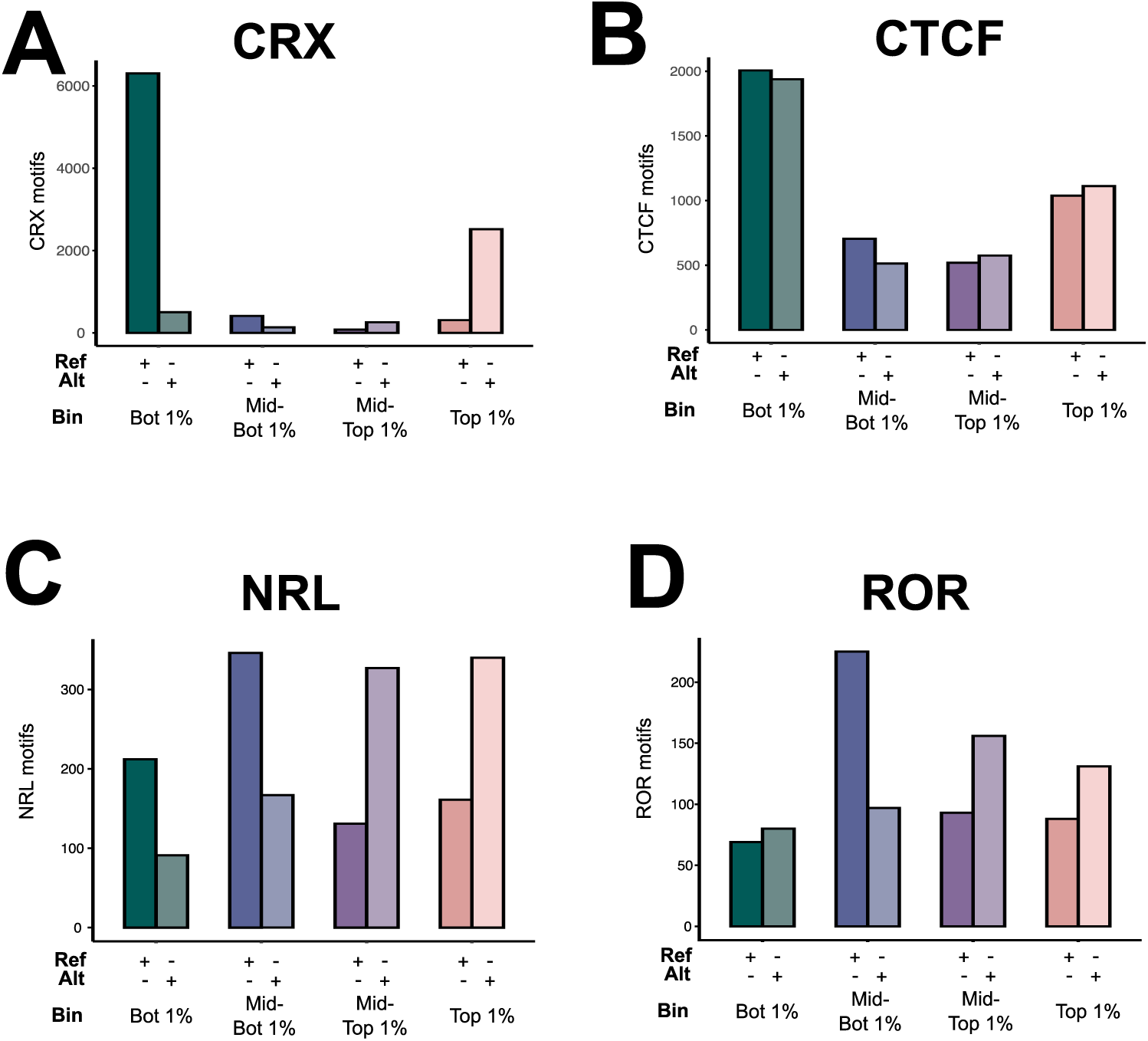
Numbers of motifs as scored by FIMO in reference and variant sequences for bins highlighted in Figure 3A. Motifs shown are CRX (A), CTCF (B), NRL(C), and ROR (D).

**Supplementary Figure 6.**
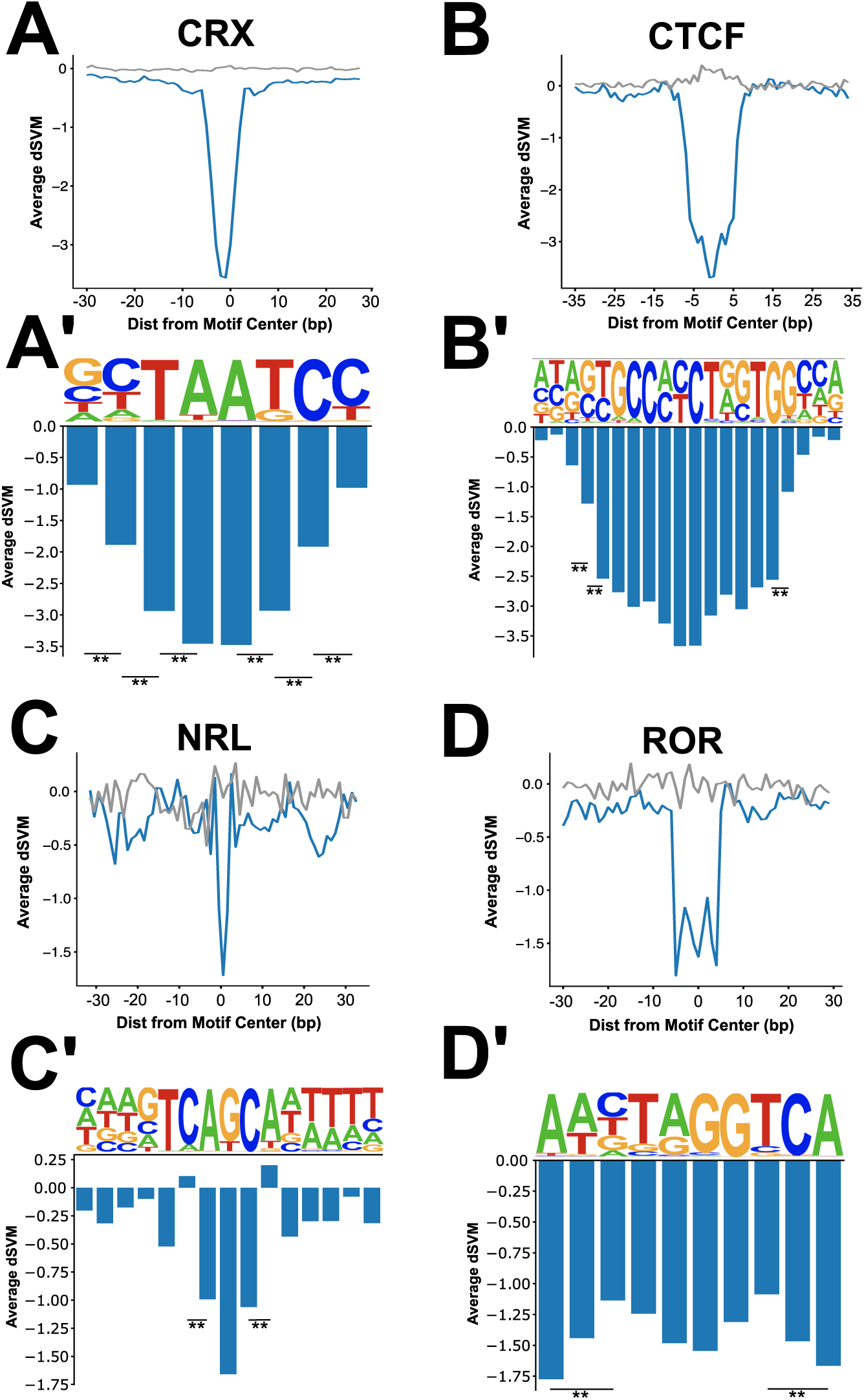
(A-D) Line plots showing the average deltaSVM for SNVs +/- 25 bp around the core motifs shown in Supplementary Figure 3. ATAC model shown in blue, shuffled model in gray. (A’-D’) Bar plots showing the average deltaSVM for SNVs on a base pair resolution within the core motifs of those shown in Supplementary Figure 3. (**: p< 0.002)

